# Benchmarking DNA binding affinity models using allele-specific transcription factor binding data

**DOI:** 10.1101/2023.12.15.571887

**Authors:** Xiaoting Li, Lucas A. N. Melo, Harmen J. Bussemaker

## Abstract

Transcription factors (TFs) bind to DNA in a highly sequence-specific manner. This specificity can manifest itself *in vivo* at heterozygous loci as a difference in TF occupancy between the two alleles. When applied on a genomic scale, functional genomic assays such as ChIP-seq typically lack the statistical power to detect allele-specific binding (ASB) at the level of individual variants. To address this, we propose a framework for benchmarking sequence-to-affinity models for TF binding in terms of their ability to predict allelic imbalances in ChIP-seq counts. We show that a likelihood function based on an over-dispersed binomial distribution can aggregate evidence for allelic preference across the genome without requiring statistical significance for individual variants. This allows us to systematically compare predictive performance when multiple binding models for the same TF are available. We introduce PyProBound, an easily extensible reimplementation of the ProBound biophysically interpretable machine learning framework. Configuring PyProBound to explicitly account for a confounding sequence-specific bias in DNA fragmentation rate yields improved TF binding models when training on ChIP-seq data. We also show how our likelihood function can be leveraged to perform *de novo* motif discovery on the raw allele-aware ChIP-seq counts.

## BACKGROUND

Transcription factors (TF) bind to DNA and regulate transcription in a sequence-specific manner [1]. Genome-wide association studies (GWAS) have shown that the majority of significantly associated variants are located in non-coding regions and may therefore impact gene regulation [2]. One of the key drivers of phenotypic variation is variable TF-DNA binding [3], which is commonly caused by the disruption of TF binding sites by genetic variants.

The effect of genetic variation on TF binding can be investigated through an allele-specific approach, which can assess allelic differences in functional genomic readouts directly at heterozygous loci, thus better controlling for environmental differences between individuals and cell types. For instance, allele-specific binding (ASB) can be detected as a statistically significant imbalance between the number of mapped ChIP-seq reads containing the respective alleles of a single-nucleotide variant (SNV) [4]. AlleleDB [5] is a resource that provides ASB annotations based on the 1000 Genome Project.

Allelic preference at single nucleotide variants (SNVs) may arise from the alteration of TF binding sites and therefore of TF occupancy [6]. In support of this, the AlleleDB [5] and ADASTRA [7] studies both reported concordance between ASB calls and motif disruption. However, an important fraction of ASB instances cannot be explained in terms of direct alteration of TF binding affinity [8, 9]. In these cases, the observed allelic imbalance may be due to variation in indirect TF binding or chromatin accessibility. Prediction of SNV effects on TF binding is also limited by the quality of the TF binding models, especially for weaker sites [10].

Quantitative knowledge about a TF’s ability to bind to specific DNA sequences is essential to understanding its function. TF binding models or “motifs” can be used to identify potential binding sites by scanning DNA sequences of cis-regulatory regions such as promoters and enhancers [1]. New techniques for probing TF-DNA interactions have greatly expanded our knowledge of TF binding specificity. High-throughput *in vitro* assays can readily characterize intrinsic TF binding preferences on a large scale. SELEX (systematic evolution of ligands by exponential enrichment) is one of the most widely used *in vitro* assays, assessing binding of a purified TF protein across a large pool of random DNA ligands through affinity-based selection over multiple rounds [11]. High-throughput SELEX data are available for hundreds of TFs [12].

High-throughput *in vivo* assays study TF binding under a specific biological condition in a particular cell type, and may reflect more complex cooperative TF binding with co-factors also present in the nucleus. Chromatin immunoprecipitation (ChIP) assays, in which proteins are crosslinked to their DNA binding sites before fragmentation and immunoprecipitation, are among the most widely used methods for probing the *in vivo* occupancy of TFs, and can be coupled with next-generation DNA sequencing (ChIP-seq) [13]. Thanks to efforts such as ENCODE, ChIP-seq data are also available for hundreds of TFs [14, 15]. Alternative *in vivo* assays for probing TF binding include ChIP-exo, which adds an exonuclease digestion step [16], and CUT&Tag [17], which does not require crosslinking.

Many computational studies have attempted to improve the prediction of genetic variant effects on TF binding. ProBound, a flexible machine learning framework recently developed in our lab [18] directly fits a biophysical model to multiple rounds of SELEX data, while accounting for non-specific binding, dependencies between nucleotide positions, and multiple binding modes. ProBound can systematically and consistently analyze data from different types of SELEX experiments, and is capable of identifying low-affinity sites and capturing the impact of co-factors and DNA methylation. DNA binding models learned from high-throughput SELEX data using ProBound generally outperform binding motifs from other resources, including JASPAR [19], DeepBind [20], and HOCOMOCO [21] when predicting *in vivo* DNA occupancy [18]. This suggested that ProBound may also have great potential for predicting the impact of genetic variation on TF-DNA binding, which is the topic of the present study.

Our previous study [18] included proof-of-concept that ProBound can be used to analyze single-end ChIP-seq data in a way that avoids the statistical distortion associated with peak calling. A DNA binding model inferred from ChIP-seq data for the glucocorticoid receptor was quantitatively consistent with a model derived from SELEX data for the same TF. The original implementation of ProBound however was not designed to able to handle DNA libraries with variable sequence lengths, which precluded optimal analysis of paired-end ChIP-seq data.

For this study, we created PyProBound, a machine learning framework based on PyTorch that reimplements the methodology of [18] in a more flexible and modular manner, and at the same time is fully backwards compatible with the original Java implementation of ProBound. We use PyProBound to learn TF binding models from paired-end ChIP-seq, ChIP-exo, and CUT&Tag data without the need for any peak calling. It is in fact not even necessary to map any reads to the genome if the DNA fragments defined by the read pairs are long enough. We do find that our peak-free approach requires explicit modeling of the local sequence dependence of DNA fragmentation rate, but this is straightforward to implement in the PyProBound framework.

Using the widely studied human transcription factor CTCF as an illustrative example, we compare sequence-to-affinity models derived from various types of TF binding data in terms of their ability to predict the impact of genetic variation on TF binding. To this end, we construct a likelihood function that can quantify, on a genome-wide scale, to what extent allelic preference can be explained from DNA sequence alone. It does so without the need to make any calls of ASB at the level of individual variants. We show that the same genome-wide likelihood function can be leveraged to perform *de novo* motif discovery directly on allele-aware binding data using PyProBound. Taken together, our results underscore both the usefulness of resources such as AlleleDB and the extensibility of PyProBound.

## RESULTS

### Prediction of CTCF allele-specific binding events using sequence-to-affinity models

To assess to what extent sequence-to-affinity models can predict allele-specific binding effects, we used human ASB annotations from AlleleDB [5] and predicted the SNV’s effect on TF binding affinity. We chose to focus on the insulator protein CTCF due to the abundance of variants (2,231 SNVs) with significant evidence of ASB for this factor. The MotifCentral database contains hundreds of ProBound models trained on *in vitro* binding data from HT-SELEX [12] and SMiLE-seq [15] assays. We used the MotifCentral model for CTCF to predict allele specific binding at heterozygous SNV loci that were previously found to have significant allelic bias in ChIP-seq coverage (**Figure 1A**). For each SNV observed to have ASB, we predicted cumulative CTCF binding affinity from the DNA sequence around the SNV by summing over all offsets of the TF-DNA binding interface relative to the SNV, separately for the reference and the alternative allele (**Figure 1B**; see **Methods** for details).

**Figure 1:**
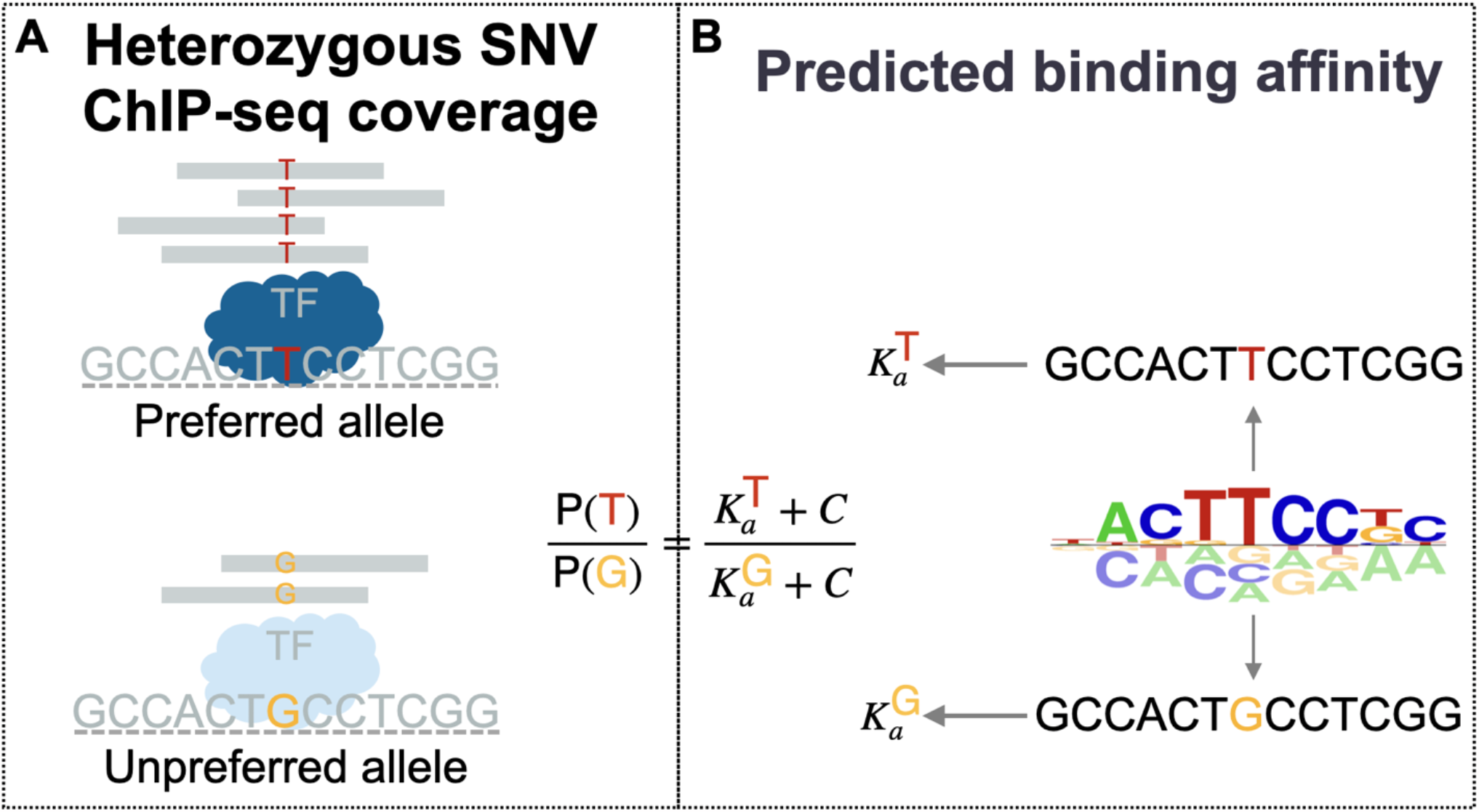
Overview of the allele-specific binding data and binding affinity scoring. (**A**) For each SNV, the preferred and unpreferred allele are defined in terms of ChIP-seq read coverage. (**B**) The binding affinity for each SNV is computed as the sum of relative affinity scores over all possible offsets.

A key question is to what extent the direction of the *in vivo* allelic imbalance in binding is in concordance with the difference in the binding affinity as predicted by the sequence-to-affinity model. Encouragingly, the SELEX-derived model for CTCF in MotifCentral was able to predict the direction of ASB for CTCF binding sites covering a 1,000-fold range in predicted affinity (**Figure 2**). Only two other TFs represented in AlleleDB have at least 100 SNVs with evidence for ASB (387 SNVs for PU.1 and 189 SNVs for EBF1). While there is still a trend that the favored allele corresponds to stronger TF binding predicted by the MotifCentral model for these factors (**Supplemental Figure S1**), the coverage of individual SNVs by ChIP-seq reads is far too low to detect ASB across the genome. This underscores the value of sequence-based prediction of ASB, or even better, quantifying difference in binding affinity between the two alleles, provided that it is possible to assess the reliability of such predictions.

**Figure 2:**
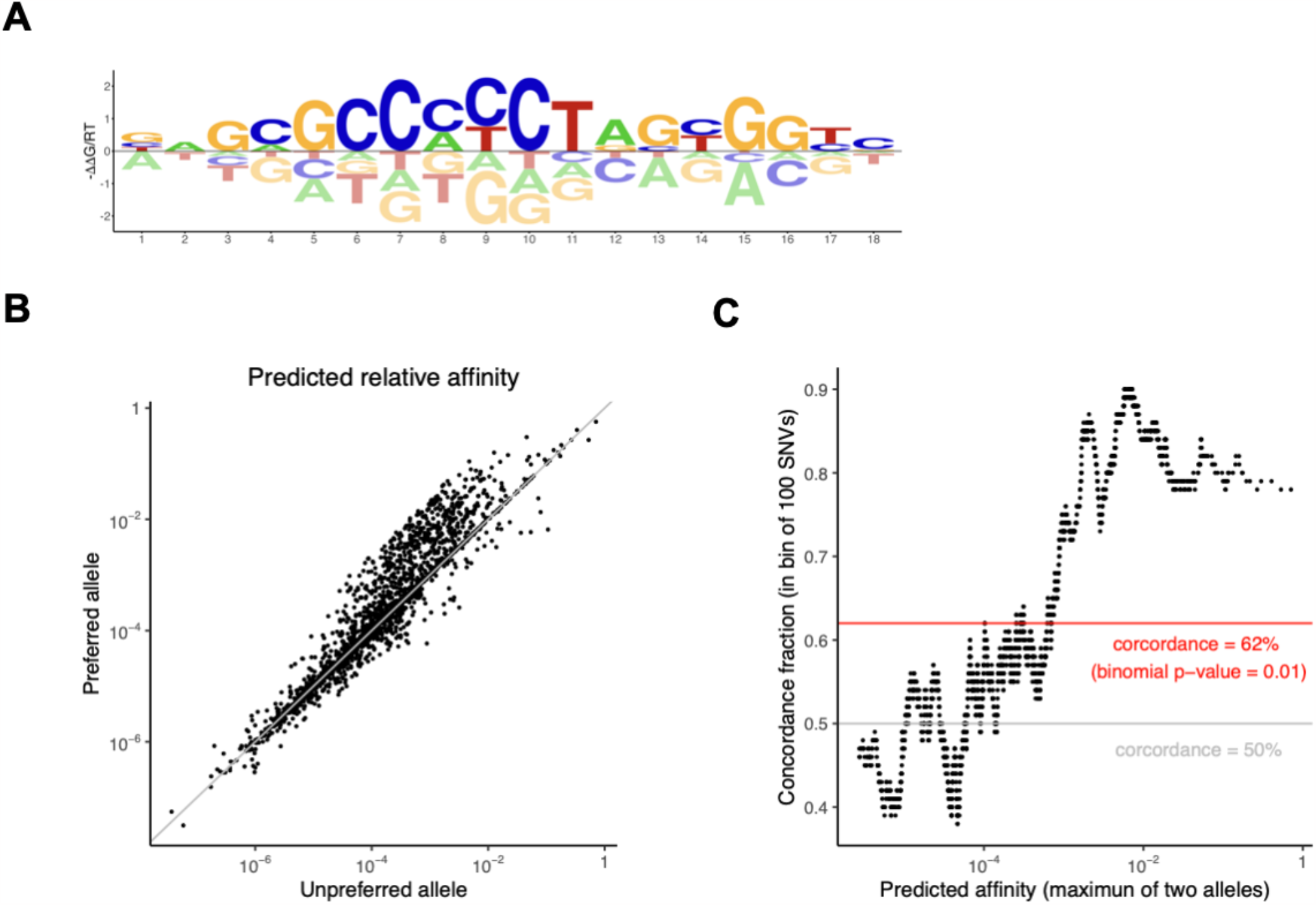
Predicting allele-specific binding by CTCF using a sequence-to-affinity model. (**A**) Energy logo [26] representation of a CTCF binding model from MotifCentral capable of making accurate predictions of binding affinity. (**B**) Comparison of predicted affinity between the preferred allele (i.e., the one covered by a significant majority of the ChIP-seq reads) and the unpreferred allele. Each dot corresponds to a distinct single-nucleotide variant (SNV) at which significant ASB was detected in the AlleleDB study [5]. (**C**) Concordance of allelic preference as (i) predicted from DNA sequence and (ii) observed using ChIP-seq. The x-value corresponds to the greater of the predicted affinity of the two alleles; the y-value is the concordance ratio in bins of 100 consecutive SNVs as ranked by predicted affinity.

### A metric for binding model quality based on observed allelic preference across the genome

Since for most TFs the fraction of SNVs that are associated with allele-specific binding on an individual basis is very small, we set out to find a way to aggregate below-threshold evidence for allelic preference across the genome. We settled on a likelihood framework that uses the same beta-binomial distribution used by [5] to model allelic counts for AlleleDB; however, rather using this model to reject the null hypothesis that both alleles are equally probable, we used the likelihood function to quantify the performance of sequence-based predictors of allelic preference on a genome-wide scale (see **Methods** for details). Importantly, this was done without regard to the statistical significance of allelic imbalance at the level of individual SNVs.

AlleleDB [5] used a beta-binomial test with a probability of 1/2 for both alleles to detect instances of ASB from allele-aware ChIP-seq data. By contrast, we here used allele-specific ChIP-seq counts aggregated across multiple individuals, and re-estimated the overdispersion parameter underlying the distribution by maximizing the likelihood function. We reasoned that using SNV-specific allelic probabilities based on CTCF binding affinities predicted from DNA sequence in the beta-binomial model should improve the likelihood, compared to a model with a fixed probability of 1/2 as an equal-preference control.

We found that this was indeed the case for the sequence-to-affinity model for CTCF in MotifCentral. To assess the statistical significance of the difference, we used bootstrapping to sample the log-likelihood distribution for each binding model separately (**Figure 3A**; see **Methods** for details). Compared to the equal-preference control, the MotifCentral model has a significantly higher mean log-likelihood (LL = – 3.2021 for control; LL = –3.2356 for MotifCentral; Wilcoxon test p < 10^−8^). Consistently, the maximum-likelihood estimate of the overdispersion parameter is significantly lower (0.0916 for control; 0.0990 for MotifCentral), indicating superior performance in explaining allelic differences in ChIP-seq counts. As expected, including all offsets at which the binding model overlaps with the SNV are important when predicting cumulative affinities from its flanking DNA sequence (**Supplemental Figure S2**); we saw more modest further improvement in the likelihood when also including offsets that do not overlap with the SNV, which still have the potential to contribute to the ChIP enrichment.

**Figure 3:**
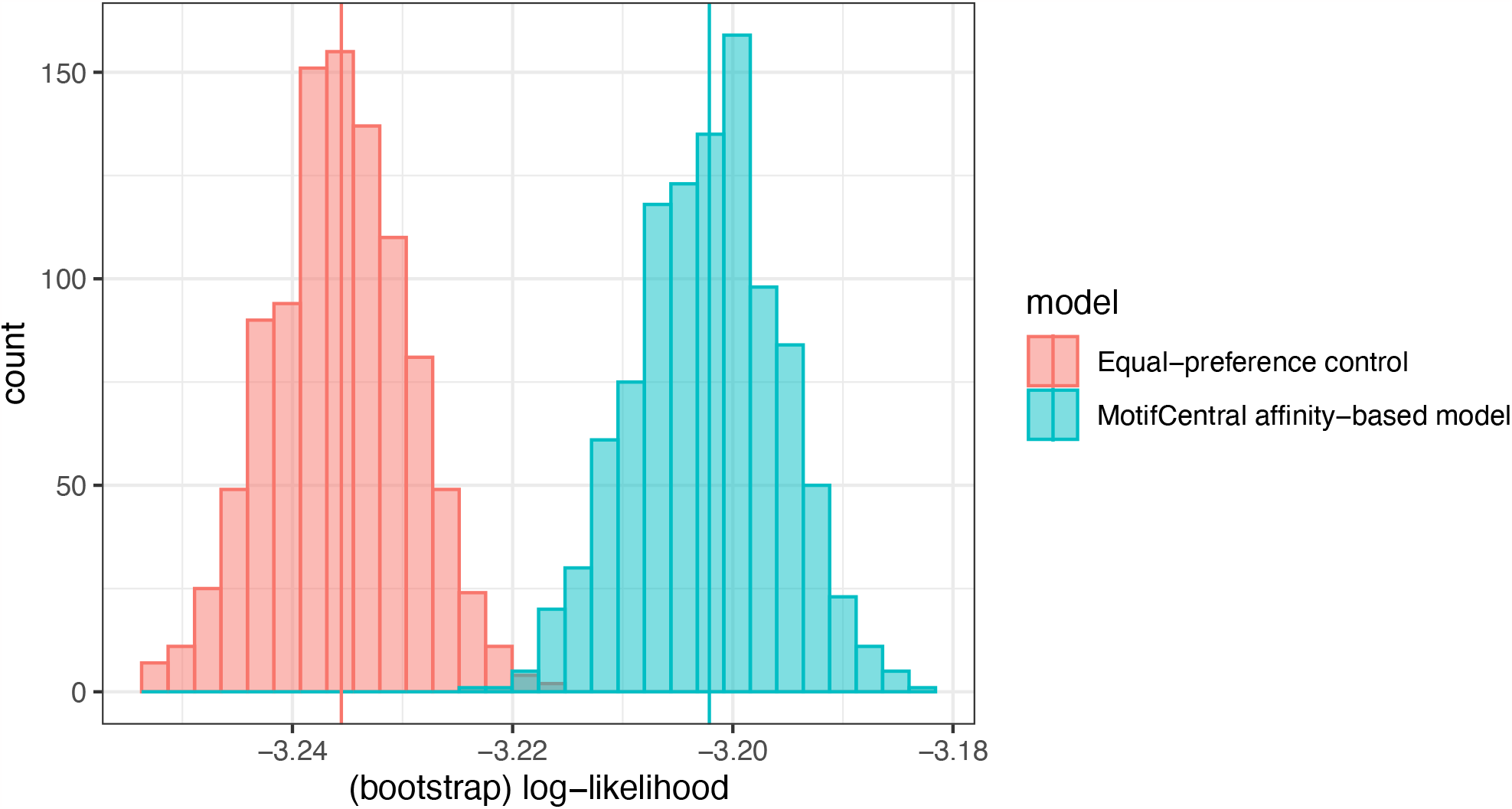
Bootstrap distributions of log-likelihood for CTCF. The histograms show the bootstrap distributions of log-likelihood for 1000 resamples, based on the affinity-based likelihood model (allelic ratio predicted from genome sequence using binding model) or the control model (which assumes the alleles are equiprobable). The vertical line indicates the observed value of log-likelihood from each model.

We applied the same analysis to PU.1 and EBF1, and again the MotifCentral model showed statistically significant predictive performance when aggregating evidence across all SNVs (**Supplemental Figure S3**). Overall, our likelihood-based framework for integrating evidence across the genome based on the beta-binomial distribution provides a way to quantitatively compare sequence-based predictors of allelic preference in an aggregate manner that is not limited by the number of SNVs that reaches statistical significance for calling ASB at the individual-locus level.

### Taking DNA fragmentation bias into account when analyzing ChIP-seq data yields superior sequence-to-affinity models

So far, the TF binding models we used to score binding affinity were all derived from *in vitro* SELEX data. We wondered whether models derived in an allele-agnostic manner from *in vivo* assays such as ChIP-seq and CUT&Tag would be better at explaining the allelic imbalances reflected in the allele-aware ChIP-seq data. To this end, we configured PyProBound to analyze paired-end *in vivo* data without peak calling (see **Methods** for details). Notably, when using the ChIP-seq data for CTCF from ENCODE, we found that the length of the merged read pairs was large enough for PyProBound to infer accurate binding models without any mapping to the genome.

Initially, when using PyProBound to fit a single binding mode intended to represent the DNA binding specificity of CTCF (**Supplemental Figure S4A**), the associated positional profile also learned by PyProBound indicated a strong bias at the ends of the ChIP-seq fragments (**Supplemental Figure S4B**). We interpreted this as confounding between the sequence preferences of CTCF binding and an unknown local sequence dependence of DNA fragmentation at the ends of the paired reads. To address this, we next configured PyProBound to fit a model that accounts for two multiplicative effects simultaneously: (i) the sequence dependence of the rate of DNA fragmentation during sonication at the two observed ends of the fragment; and (ii) the sequence dependence of CTCF binding, which determines the probability that the fragment is crosslinked with this TF during immunoprecipitation (see **Methods** for details). This more sophisticated use of PyProBound when learning a sequence-to-affinity model from allele-agnostic ChIP-seq for CTCF led to significantly improved performance when predicting allelic preference across the genome (**Supplemental Figure S4C**) and made the positional bias disappear (**Supplemental Figure S4D**). A simpler approach, in which we truncated each fragment to 200bp relative to its center, which due to the varying length of the fragments obscures the fragmentation bias, did not perform equally well in terms of predicting enrichment during the immunoprecipitation step (**Supplemental Figure S4E, F**).

To summarize the performance of the many different sequence-to-affinity models available for CTCF, we again used our likelihood-based metric. The results were consistent with expectation in various regards: Models trained on *in vivo* binding data outperformed those trained on *in vitro* data (**Figure 4**). Additionally, ChIP-seq-derived models trained with fragmentation bias modeling outperformed those that truncate all fragments to the same length (LL = –3.2086 for length truncated; LL = –3.1995 for fragmentation modeling; Wilcoxon test p < 10^−8^). Moreover, the PyProBound model inferred from CUT&Tag data has better performance than either the PyProBound model inferred from ChIP-seq or ChIP-exo data for CTCF or motif models for CTCF from other resources (**Figure 4**).

**Figure 4:**
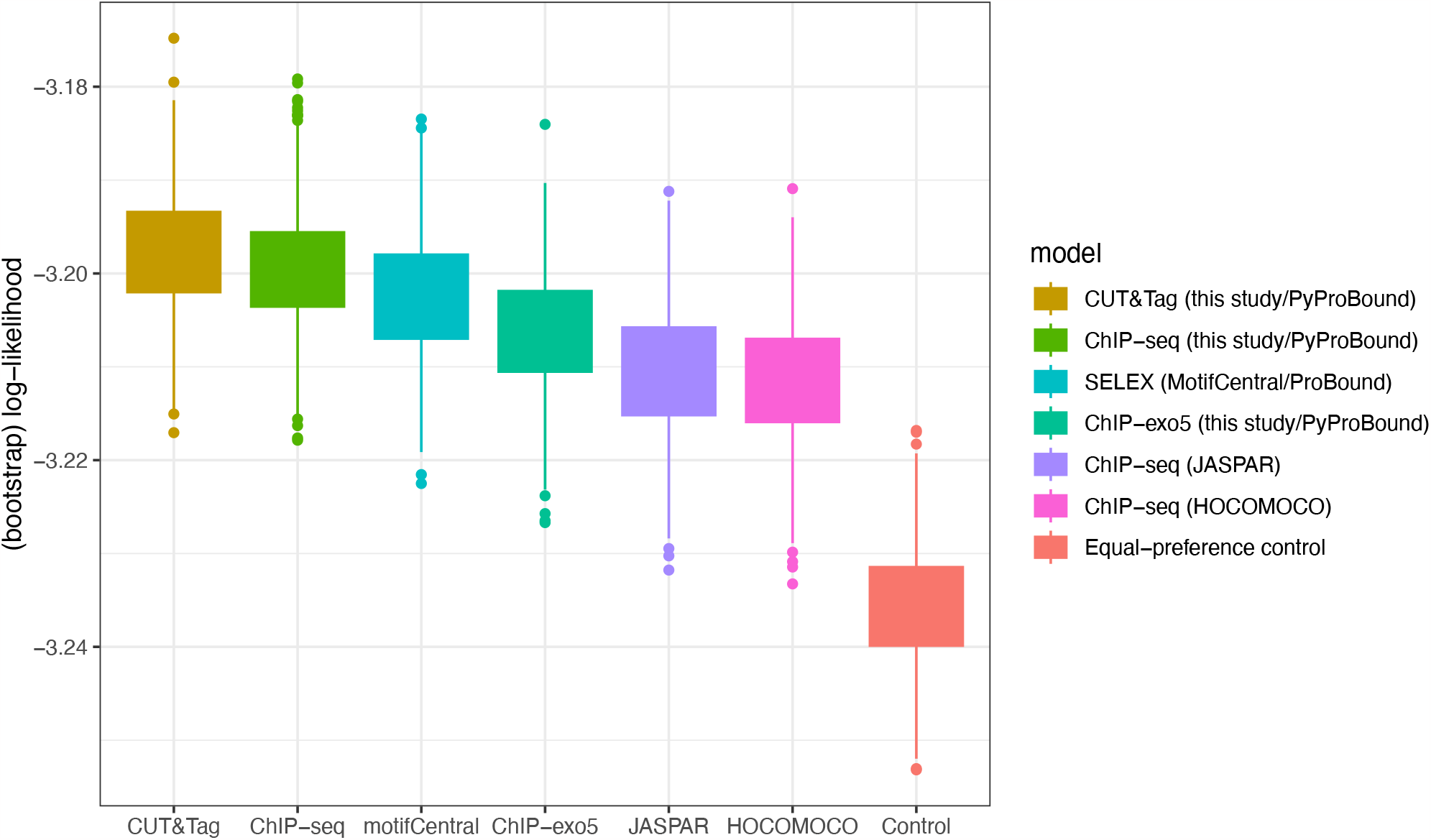
Comparison between bootstrap distributions of log-likelihood across various models for CTCF binding. The boxplot shows the comparison in log-likelihood between the control model and affinity-based models based on different CTCF binding motifs, including a CUT&Tag-derived model introduced in this study and motifs and motifs from various resources.

### *De novo* motif discovery from allele-aware ChIP-seq counts

Our success in predicting allelic preference on a genome-wide scale using independently derived TF binding specificity models suggest that the direct modulation of TF binding by SNVs can explain allele imbalance to a significant degree. We therefore wondered whether the same beta-binomial likelihood function could be leveraged to perform *de novo* motif discovery purely by trying to explain allelic imbalances in ChIP-seq counts from DNA sequence, while taking variation in combined ChIP-seq coverage among SNVs for granted. To our knowledge, such an approach, which controls for variation in chromatin context in a unique way by assuming that the local molecular environment of a given SNV on the respective homologs of the chromosome on which it resides, has not been explored before. We configured ProBound to optimize the beta-binomial likelihood function not only with respect to the overdispersion parameter and non-specific binding coefficient as was done above, but also with respect to the position-specific free-energy parameters of the TF binding model (see **Methods**). Since we could not compare the resulting model on the benchmark data it was trained on, we instead compared its energetic parameters against previously published models. The ASB-derived model showed excellent agreement with the MotifCentral model (Pearson r = 0.94; **Figure 5**), indicating that CTCF allelic imbalance is indeed driven by direct alteration of sequence-specific CTCF binding.

**Figure 5:**
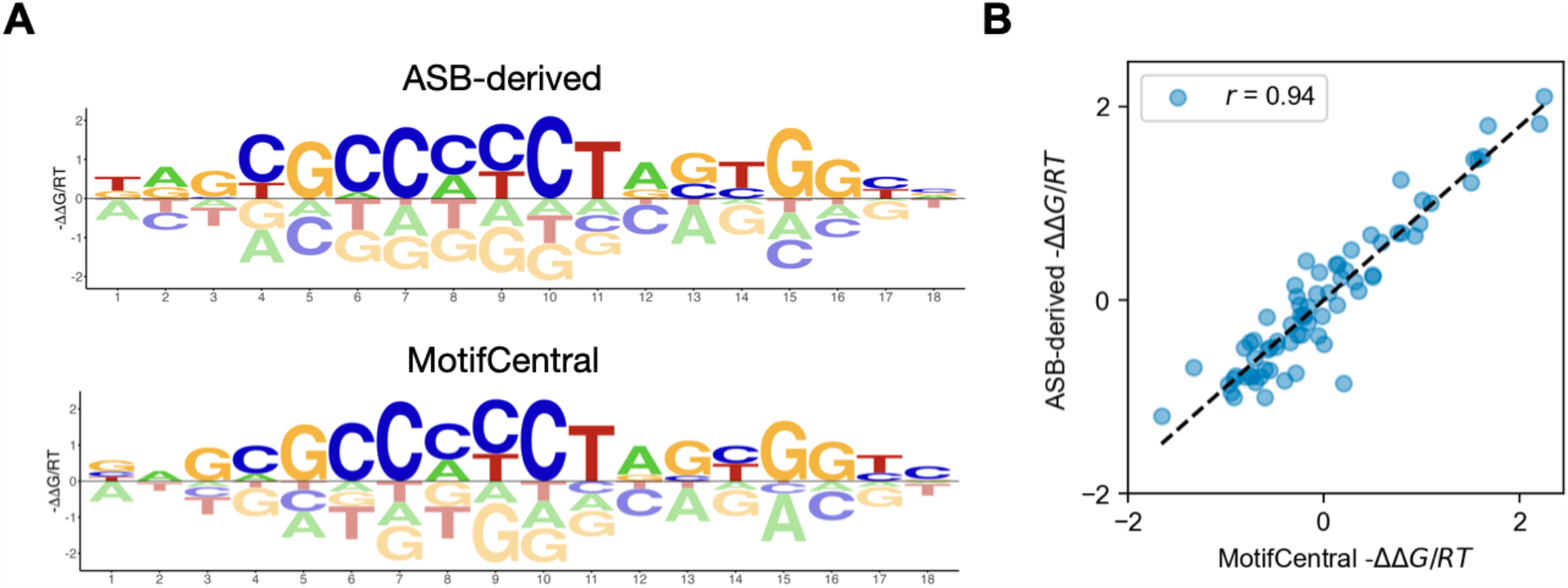
Comparison of *de novo* ASB-derived and existing MotifCentral models. (**A**) Energy logos for binding model inferred from allele-aware ChIP-seq data from [5] using PyProBound, and from SELEX data using the original version of ProBound [18]. (**B**) Direct comparison of free energy parameters in the respective models, with each point corresponding a unique base/position combination in the logo.

## DISCUSSION

In this study, we used allele-aware ChIP-seq data from AlleleDB [5] to assess the performance of DNA binding models when predicting allelic preference using sequence-based scoring of variation in binding affinity. The allele-specific approach is useful for assessing the prediction of genetic effects on *in vivo* TF binding. Leveraging the availability of many types of *in vitro* and *in vivo* binding data for the human transcription factor CTCF, we demonstrated that CTCF binding affinity as predicted using binding models trained with the ProBound algorithm is in concordance with allelic imbalance observed in allele-aware ChIP-seq data. This underscores the potential of ProBound for predicting SNV effects from DNA sequence alone.

We developed a likelihood framework based on the (over-dispersed) binomial distribution that can be used to quantify in an unbiased manner how well the predicted binding affinities can explain allelic preference on a genome-wide scale. This makes it possible to leverage the large number of SNVs for which allele-aware ChIP-data do not provide enough statistical power to make calls of ASB on an individual-SNV basis. In fact, for most TFs, it is not possible to detect any instances of ASB using per-SNV analysis of ChIP-seq counts. We propose that our method for aggregating evidence for allelic preference across the genome can be broadly useful for benchmarking sequence-based predictors of TF binding affinity across many different cellular contexts and many different TFs.

As expected, when comparing the performance of various binding models for CTCF when predicting allelic differences in TF binding as assayed *in vivo* using ChIP-seq, ChIP-exo, or CUT&Tag, models derived from *in vivo* data in an allele-agnostic manner generally perform better than those derived from *in vitro* binding data, even when the role of co-factors is not explicitly considered. In particular, CUT&Tag-derived models outperform ChIP-based approaches, perhaps by avoiding sequence-specific biases due to crosslinking. We also showed that our genome-wide likelihood function can be easily leveraged to perform *de novo* motif discovery from allele-aware ChIP-seq data, thanks to the flexible nature of our PyProBound software.

Taken together, our findings underscore the value of optimal methodology for estimation the binding energy parameters, which was our overall motivation for developing ProBound and related tools [18, 22]. ProBound already naturally accounts for biases in the ChIP input library, by only focusing on explaining how the immunoprecipitation step leads to further enrichment. Therefore, this bias is distinct from the DNA fragmentation bias that we explicitly account for with PyProBound using the technique described in this paper.

## CONCLUSIONS

It is an exciting prospect that for a future generation of more elaborate TF binding models that explicitly account for cooperative interactions among sets of interacting TFs, some of which may be inferred using the ProBound algorithm [18], our likelihood framework could again provide an objective an unbiased metric for assessing and comparing their predictive performance in an *in vivo* context. This includes accounting for cell-type-specific TF interactions, as long as allele-aware TF binding data in a matching cell line or tissue would be available to define a benchmark metric.

## METHODS

### Collection of Allele-Specific Binding SNVs

Allele-specific TF binding data was downloaded from the AlleleDB database [5], which contains allele-specific annotations for the 1000 Genomes variant catalogue (https://doi.org/10.1038/nature15393). The AlleleDB authors reprocessed hundreds of ChIP-seq assays on tissue samples from 14 human individuals, mapping reads to a personal genome, and identifying allelic imbalance using a beta-binomial test. The SNVs classified as ‘accessible’ in AlleleDB comprises heterozygous loci that have at least the minimum number of reads needed to be statistically detectable, and consists of both SNVs with statistically significant ASB and non-ASB SNVs that can serve as controls. In the present study, we used the raw ChIP-seq counts for the reference and alternative allele as input to our modeling. We primarily examined CTCF in our analyses due to its relative abundance of ASB (2,231 ASB SNVs, 44,422 control SNVs). We also analyzed data for EBF1 (189 ASB SNVs) and SPI1 (387 ASB SNVs).

### Sequence-based prediction of protein binding affinity for ASB prediction

A TF can bind genomic DNA near a given SNV at various offsets and in either orientation, so we computed the cumulative binding affinity for each allele of the SNV as a “sliding-window” sum of relative binding affinity values as predicted by a sequence-to-affinity model. We used the R language and the BSgenome.Hsapiens.UCSC.hg19 package from Bioconductor.org to construct reference and alternative DNA sequences adding 29 bases of flanking sequence on each side of the SNV, sufficient to accommodate binding models of various sizes.

### TF-DNA binding models used in the analyses

We downloaded TF binding models derived from HT-SELEX data using ProBound [18] from motifcentral.org. In addition, we collected TF binding motifs for CTCF, SPI1 and EBF1 from HOCOMOCO [21], and JASPAR [19]. We scored DNA sequences using the ProBound scoring tool (probound.bussemakerlab.org).

### *In vivo* binding data used for de novo motif discovery using ProBound

Raw FASTQ files corresponding to the paired-end CTCF ChIP-seq were downloaded from ENCODE (encodeproject.org) using accession numbers ENCLB048DBS and ENCLB581JXH. Reads were pair-ended with BBMerge using default parameters [23]. Raw FASTQ files corresponding to the CTCF ChIP-exo and CUT&Tag were downloaded from SRA (www.ncbi.nlm.nih.gov/sra) using accession numbers SRR6736398, SRR6736390, SRR8435051, and SRR8754587. Reads were mapped with BBMap [24] using default parameters and then quality filtered using samtools view [25] with parameters `-q 30 -F 1804 -f 2` before extracting the reads.

### Beta-binomial model of genome-wide allelic effects on the binding affinity

To quantify how well the predicted binding affinity can explain the genome-wide ASB effects, we build a generalized linear model based on the beta-binomial distribution:

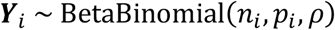

Here 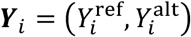 denotes raw ChIP-seq counts Y for a SNV with either reference or alternative alleles. In the beta-binomial distribution, 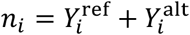 plays the role of the sample size, and ρ is the over-dispersion parameter. The binomial success rate *p*_i_ was modeled as the modified ratio of relative affinities for reference allele and alternative allele as follows:

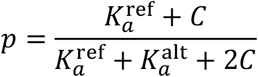

Here 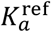 and 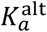 are the cumulative affinities computed as described above; C represents background binding due to indirect effects or binding of other TFs. The parameters C and ρ were estimated by likelihood maximization using the R language. The likelihood was computed by the probability density function of the beta-binomial distribution within the R package, VGAM 1.1-5 (Vector Generalized Linear and Additive Models, n.d.). The mean log-likelihood across SNVs was then computed with optimal parameters const and ρ. For the control model, a fixed 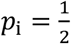 was used and the parameter ρ was estimated by likelihood maximization.

### Bootstrapping of log-likelihood

To construct the sampling distribution of log-likelihood, the SNVs were resampled 1000 times with replacement and each time the parameters of the model were estimated using the same function. The distributions of log-likelihood from 1000 bootstraps were then constructed for the control model and affinity-based model separately.

### *In vivo* binding data analysis with PyProBound

The algorithm follows the methodology published previously [18]. Briefly, the relative concentration *f*_*i,r*_ of fragment *i* in round *r* is defined in terms of a non-specific binding term α_*NS*_ and the relative binding affinity 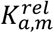 for each binding mode *m*:

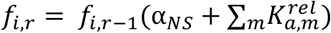

The quasi-Newton optimization method L-BFGS is used to optimize the loss function

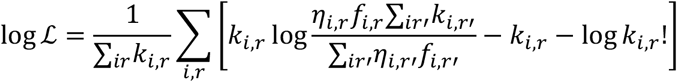

where *k*_*i,r*_ is the observed count of fragment *i* in round *r, η*_*i,r*_ is a parameter that adjusts for the read depth, and *f*_*i*,0_ = 1 by convention.

Unlike the original Java implementation of ProBound (http://github.com/RubeGroup/ProBound), PyProBound can train on sequences of varying lengths. This allows for analysis of paired-end *in vivo* binding data directly without extending or truncating reads. The enrichment of reads in the bound libraries relative to the control libraries was modeled as the product of three factors: (i) 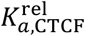 of the CTCF binding mode, summed over all sliding windows; (ii) 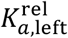 of the pair fragmentation mode scored only on the 10 base pairs in the DNA fragment; (iii) 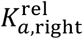 of the reverse-complemented fragmentation mode scored only on the last 10 base pairs. To each of these factors a trained non-specific binding parameter was added. Additionally, a multiplicative bias was trained for each consecutive set of five windows, to detect for bias in binding at different positions within the fragment. If the CTCF model was observed to exhibit strong bias near the ends, fragmentation modes were iteratively added until the fragmentation score of the highest-affinity sequence was lower than non-specific binding.

### PyProBound regularization

Two regularization terms were added to avoid overfitting, as published previously [18]. The first is an L2 regularization term with hyperparameter *λ* = 10^−6^ for *in vivo* models or *λ* = 10^−4^ for ASB-derived models (which required more severe regularization due to the low number of sequences trained on). The second regularization term is an exponential barrier that prevents numerical errors, and is defined as

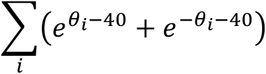

where the sum is over all parameters of the model.

## DECLARATIONS

### Ethics approval and consent to participate

Not applicable

### Consent for publication

Not applicable

### Availability of data and materials

All code for processing data files and training models can be accessed at: https://github.com/BussemakerLab/AlleleSpecificBinding

PyProBound can be downloaded at: https://github.com/BussemakerLab/PyProBound Documentation is available at: https://pyprobound.readthedocs.io

### Authors’ contributions

HJB and XL developed the ASB likelihood methodology; LANM implemented PyProBound and was responsible for its application to various data sets; XL and LANM wrote all the software and performed all analyses under the supervision of HJB; XL, LANM, and HJB wrote the manuscript.

## Acknowledgements

We thank Tuuli Lappalainen, Athena Tsu, Harshit Ghosh, H. Tomas Rube, and Chaitanya Rastogi for valuable discussions.

---

**Supplemental Figure S1:**
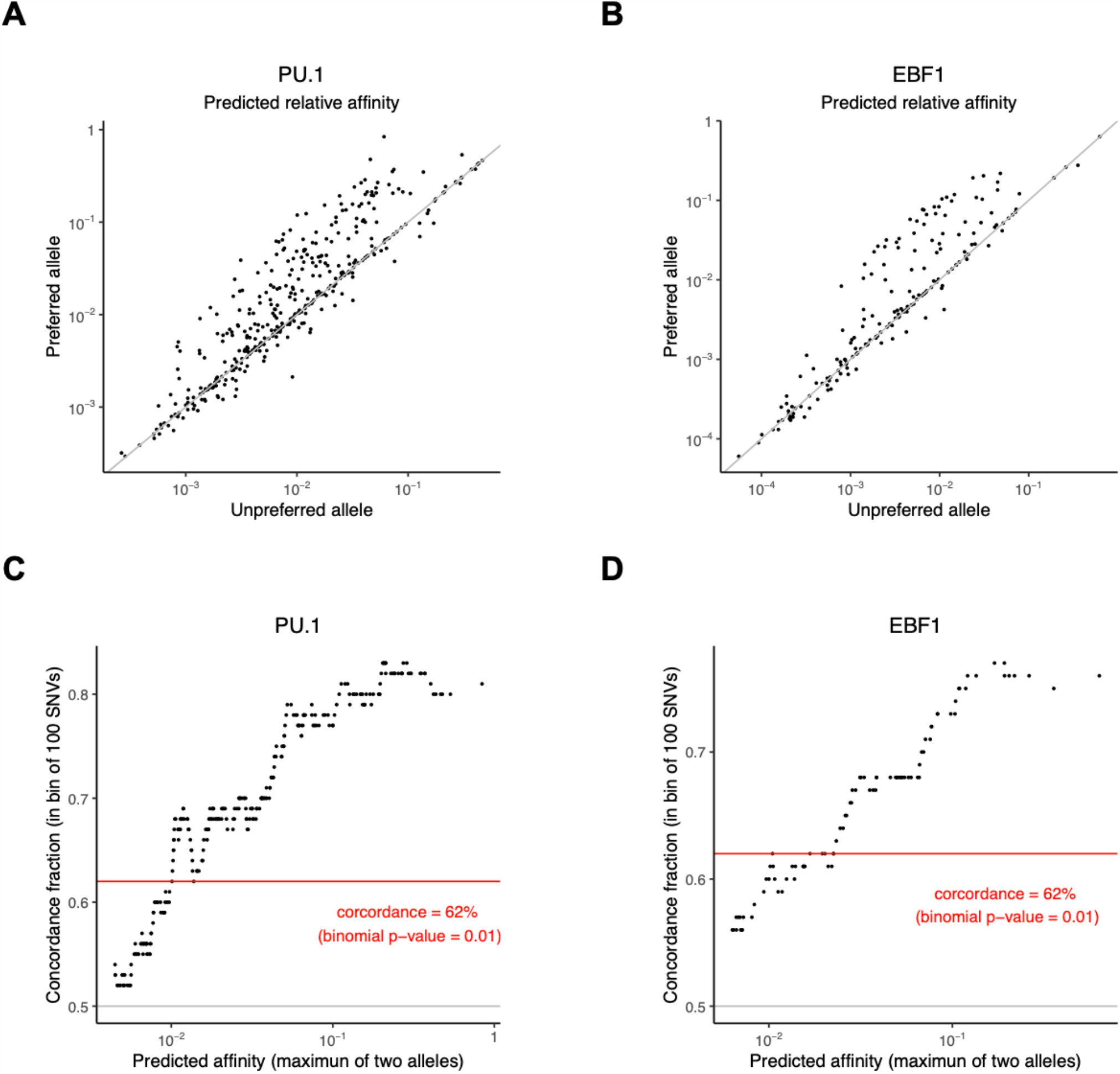
Prediction of ASB for PU.1 and EBF1 using MotifCentral models. (**A, B**) Comparison of predicted affinities between preferred alleles (the allele covered by higher ChIP-seq reads) and unpreferred alleles for PU.1 and EBF1, respectively. (**C, D**) Concordance of allelic preference as (i) predicted from DNA sequence and (ii) observed using ChIP-seq for PU.1 and EF1.

**Supplemental Figure S2:**
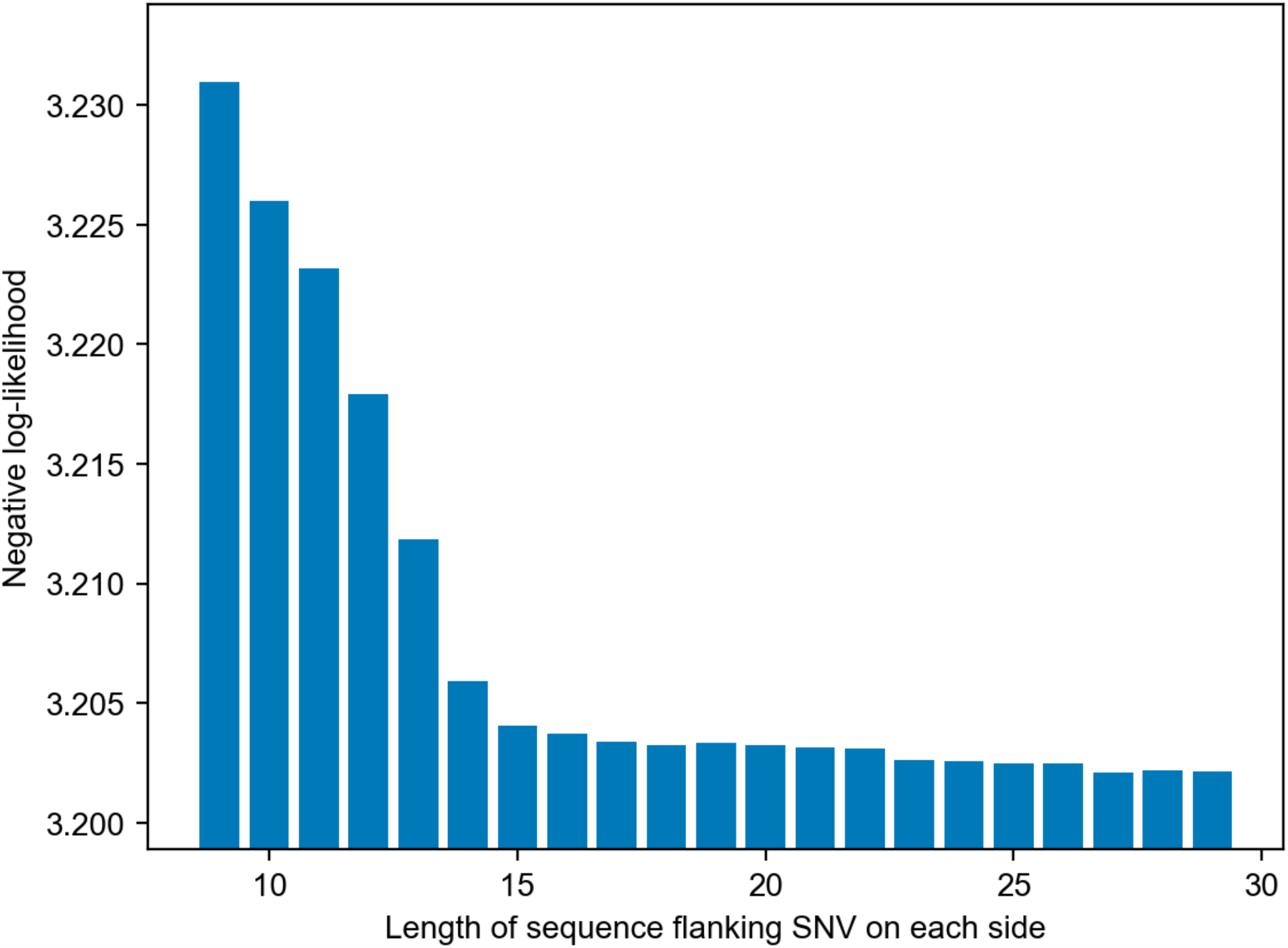
Including mode flanking data around the SNV improves the log-likelihood for beta-binomial ASB model. The plot shows the negative log-likelihood when using the MotifCentral model for CTCF.

**Supplemental Figure S3:**
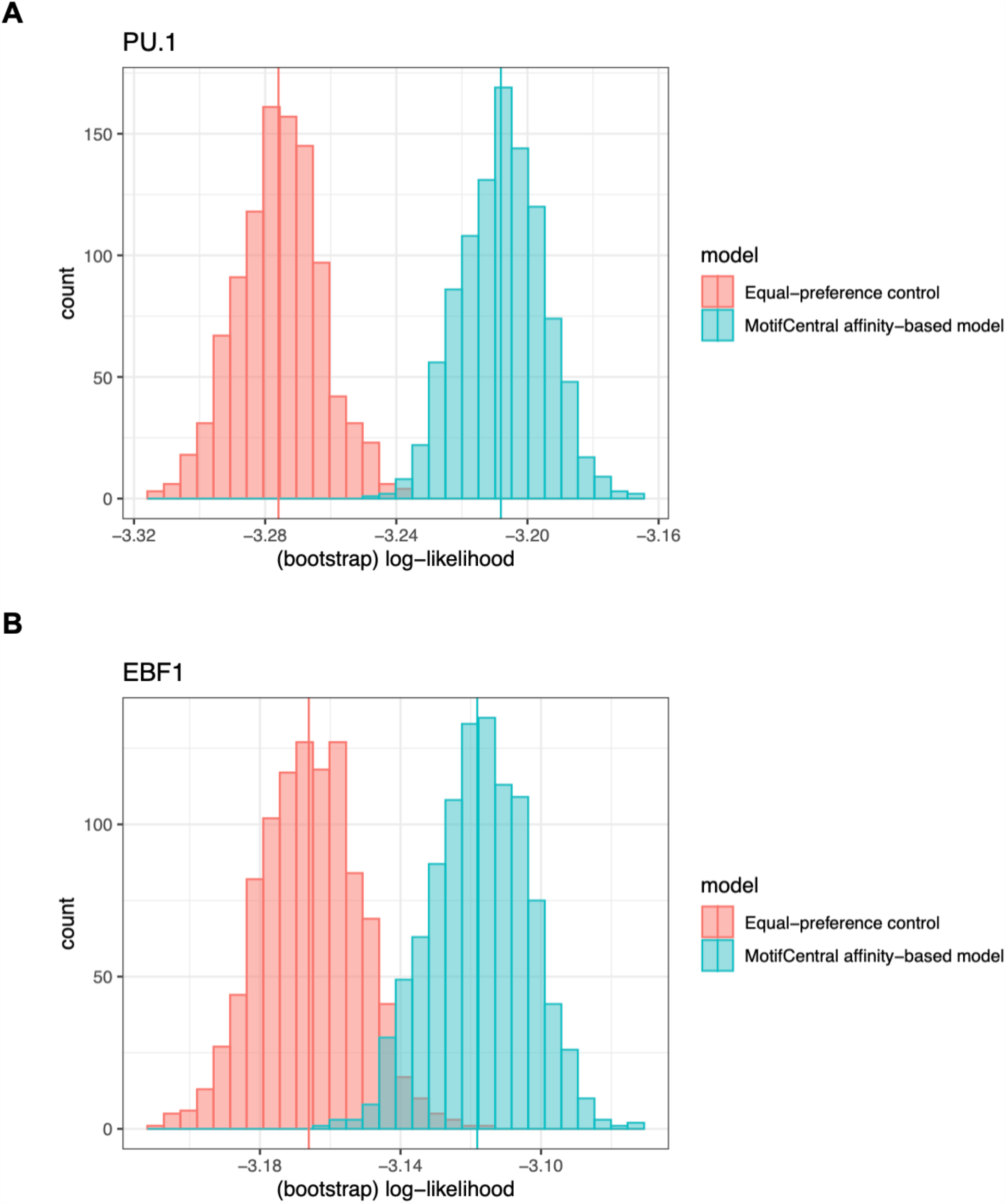
Bootstrap distributions of log-likelihood for PU.1 and EBF1. **(A, B)** Histograms showing the bootstrap distributions of log-likelihood across 1000 resamples, based on either the control model or the affinity-based likelihood model for PU.1 and EBF1.

**Supplemental Figure S4:**
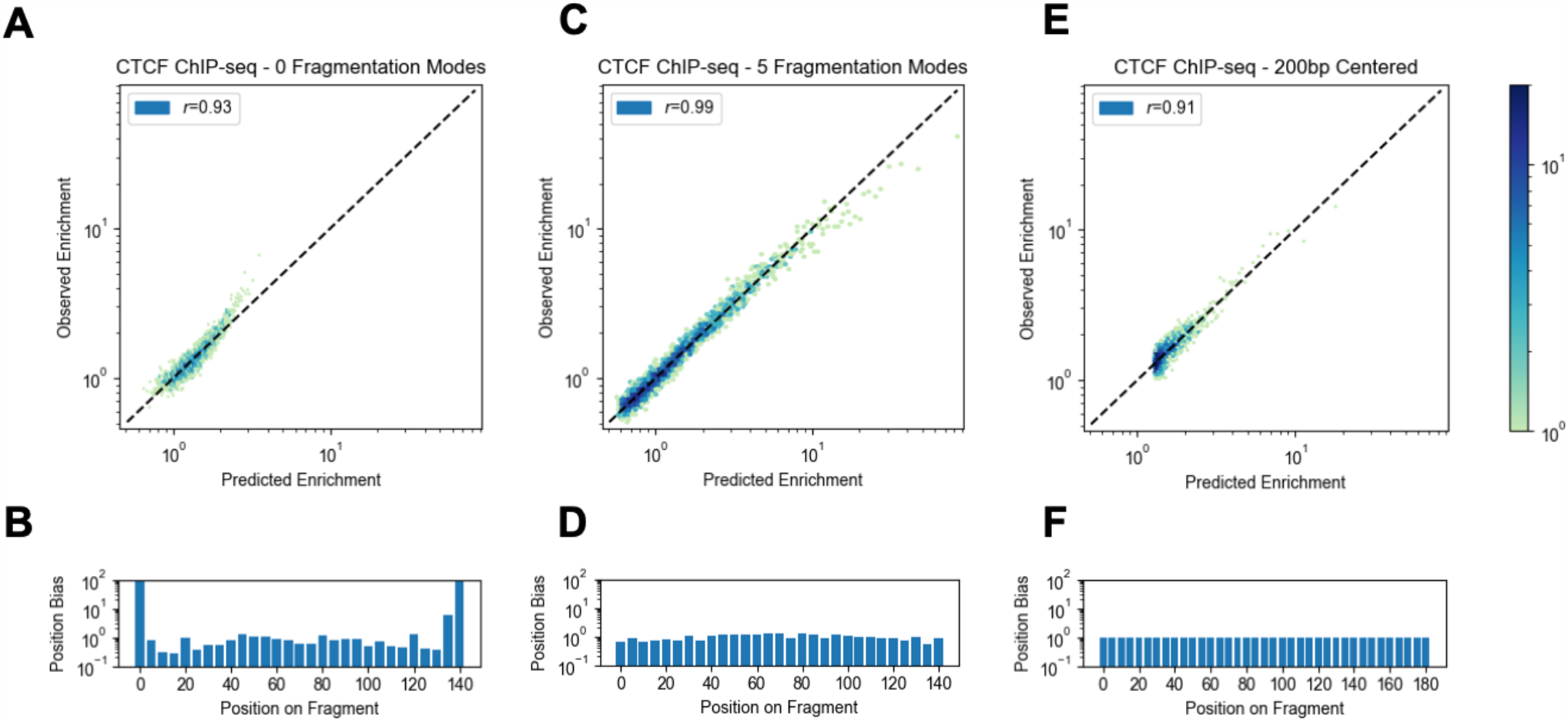
Comparison of sequence-to-affinity models inferred from *in vivo* binding data for CTCF. (**A, C, E**) Scatterplots showing predicted versus observed enrichment of DNA probes during the ChIP-seq assay. Darker colors correspond to a greater density of points, and each point corresponds to a mean over 500 sequences binned after ranking by predicted enrichment. (**B, D, E**) Positional profile relative to the IP fragment, representing the trained position bias parameter that adjusts the binding affinity at a given offset along the sequence. Compared are three different strategies for configuring PyProBound to infer a binding model from the same data for CTCF: (**A, B**) training on full-length paired-end reads corresponding to actual IP fragments that naturally vary in length; (**C, D**) training on the same data but with explicit modeling of fragmentation bias during sonication; and (**E, F**) truncation of each paired-end read to 200bp around its center.

